# Studies on Biofilm Formation by *Psuedomonas Aeruginosa* in Different Urinary Catheters

**DOI:** 10.1101/2021.02.23.432614

**Authors:** Emmanuel Asante

## Abstract

Urinary Catheters are the leading cause of healthcare –associated Urinary Tract Infections (UTIs) making their use a necessary evil. Various models and approaches had been developed to help reduce the complications associated with the use of these catheters. This study aimed at investigating how two different types of urinary catheters (Silicone Catheter and Latex Coated Catheter) support biofilm formation of *Pseudomonas aeruginosa* isolates. The growth rate of the bacteria and the rate of formation of biofilm on each urinary catheter was determined by preparing biofilm assays, ranking them using crystal violet satins and measuring their absorbance using UV-VIS spectrophotometer. A difference in the level of biofilm in the catheters were observed. The differences in the level of biofilm observed in the catheters imply that different catheters may have different susceptibility for the formation of biofilms. The study recommends the assessment of biofilm formation in the quality evaluation of catheters. Also further studies should be done to investigate how different catheter material support biofilm formation and the mechanisms involved.

## INTRODCTION

Microorganisms were first known to be planktonic, that is individual organisms living freely without any close relation. This idea was later proven to be incorrect when Van Leuwenhoek studied animalcules on his teeth, and concluded that microorganism could form close associate and live together as a whole or biofilm. (Nishitani *et al.*, 2015).

A biofilm is a group of prokaryotic microorganisms that have aggregated to form a thick layer in a form of a colony, with an inherent ability to adhere to surfaces. Biofilms can be formed on both animate and inanimate surfaces, but are mostly associated with inanimate surfaces due to the defence mechanisms propelled by living organisms against them when they found themselves on animate surfaces. The colony has a polysaccharide layer or slime layer which is porous and thus serve as a channel for nutrient transport as well as passing away waste products, it also protects the microorganism. According to the United States National Institute of Health, biofilms are responsible for 65-85% of all chronic infections. The data looks very high, but if common infections such as urinary tract infections, catheter infections and dental plaque formation are considered, it would appear this data conforms to reality. The formation of biofilm is aided by a process known as quorum sensing, the regulation of gene expression to enable microorganisms to detect and respond to cell population density. Quorum sensing is also vital for the maintenance of the already formed biofilm.

A urinary catheter is a thin, flexible tube that a person temporarily inserts into their bladder through the urethra. A catheter becomes necessary when a person has difficulties with urinating or has prostate cancer. A urinary catheter would contain droplets of urine which may form urinary stalks due to the flow of urine through it; this would inevitably become conditioned for the formation of biofilm. Although this is bound to happen, the choice of a catheter can have a significant effect on the rate of biofilm formation by chemical modifications of the polymer from which it was made. And therefore, different urinary catheters may show differences in the formation of biofilm (Hahnel *et al.*, 2015). Various models and approaches had been developed to help reduce the complications associated with the use of catheters; these approaches include the administration of antibiotics, coating catheters with antibiotics and designing new varieties in terms of material used. Despite these measures, the need for the assessment of biofilm formation for catheters, which is the major cause of urinary tract infections, has not been strongly considered. The knowledge about biofilm formation would enable us to understand why the urinary catheter is still the major cause of most urinary tract infections, regardless of the improvement in the choice of material and different varieties produced thus the need to include the assessment of biofilm formation in the quality evaluation of catheters.

*P. aeruginosa* is a gram-negative bacterium responsible for many acute and chronic diseases in plants and animals. This bacterium is an opportunistic pathogen in that it usually causes an infection during an existing disease and thus generally known to affect people whose immune system is compromised. Apart from this characteristic feature, it has the ability to form biofilm makes it difficult to be eliminated by the body defence mechanisms. The bacteria produce three polysaccharides namely aligate, Pel and Psl polysaccharides that help to maintain and strengthen the biofilm structure. Aligate is a straight unbranched polymer of D-mannuronic acid and L-guluronic acid. It aids in the structure stability of the biofilm as well as retention of nutrients whereas pel and psl are the structures formed in the initial formation of biofilm and serve as the primary structure in biofilm formation (Rasamiravaka *et al.*, 2015).

## METHODS

### Micro-organism and maintenance of the strains

*P. aeruginosa* clinical isolates were obtained and used for the experiment. The isolates were stored in small tubes and kept in the refrigerator at a very low temperature (4°C).

### Growth Conditions

Oxoid CMO405 Muller Hinton Broth was used in this experiment. A media was prepared by dissolving 2.1g of Muller Hinton broth in 100mL of distilled water. It was then autoclaved at 121°C and stored for 24 hours. A 3mL of the prepared media was transferred into three test tubes each and placed in a test tube rack. A small amount of the bacteria isolates was transferred into two test tubes leaving the last one as a control to check the sterility of the media prepared. It was left for 24 hours and observed. After, a 3mL of the media was transferred into eleven test tubes each and placed in a test tube rack. A small amount of the bacteria was inoculated into ten test tubes with a control. The absorbance of the culture was measured hourly at 600nm using a UV-VIS spectrophotometer (Spectro UV-VIS Double Beam and Scanning Auto Cell UVD-3200) and recorded. A graph was drawn to determine the growth curve of the bacterial.

### Biofilm Formation in Catheters

Two different catheters (Silicon and Latex Coated) were obtained. The catheters were cut into pieces (approximately 1cm long), placed in a chlorine solution for about ten minutes to sterilise it and then transferred using a clean forceps into an alcohol solution for about three minutes purposely to remove the bleach from the catheters. A 3mL of the media was transferred into eight test tubes each. A piece each of the catheters was transferred into four different test tubes such that four test tubes contained a piece each of one type of catheter. Three test tubes of each type of catheter were co-incubated with a small amount of the bacterial and labelled with respect to the number of days require for the measurement of the absorbance. The remaining two test tubes were used as a control; one for each type of catheter and labelled accordingly. The catheters were removed from the media and stained with crystal violet on a 24-hour interval for 3 days. The absorbance for each piece of catheter for each day was measured at 595nm using the same spectrophotometer and was recorded.

### Biofilm Formation in Microplates

Three microplates were obtained and labelled as day one, two and three. An amount of 3mL of Muller Hinton Broth was transferred into ten different holes. An amount of 0.1mL of the bacteria was inoculated in each of the ten different holes containing the Muller Hinton Broth. The media together with the inoculum was poured away and stained with crystal violet on a 24-hour interval for three days. The absorbance for each plate was measured with respect to the number of days at 595nm using a spectrophotometer and was recorded.

## RESULTS

### Growth Curve of *P. aeruginosa*

The growth curve *of P. aeruginosa* in Muller Hinton is shown in Figure1. There was no significance growth for the first two hours, an accelerated growth after the third hour, a great decline in the growth after fourth hour and a gradual increase from the fifth to seventh hour. There was a decline in the growth again from the eight to the ninth hour and a little increase after the tenth hour.

**Figure 1:**
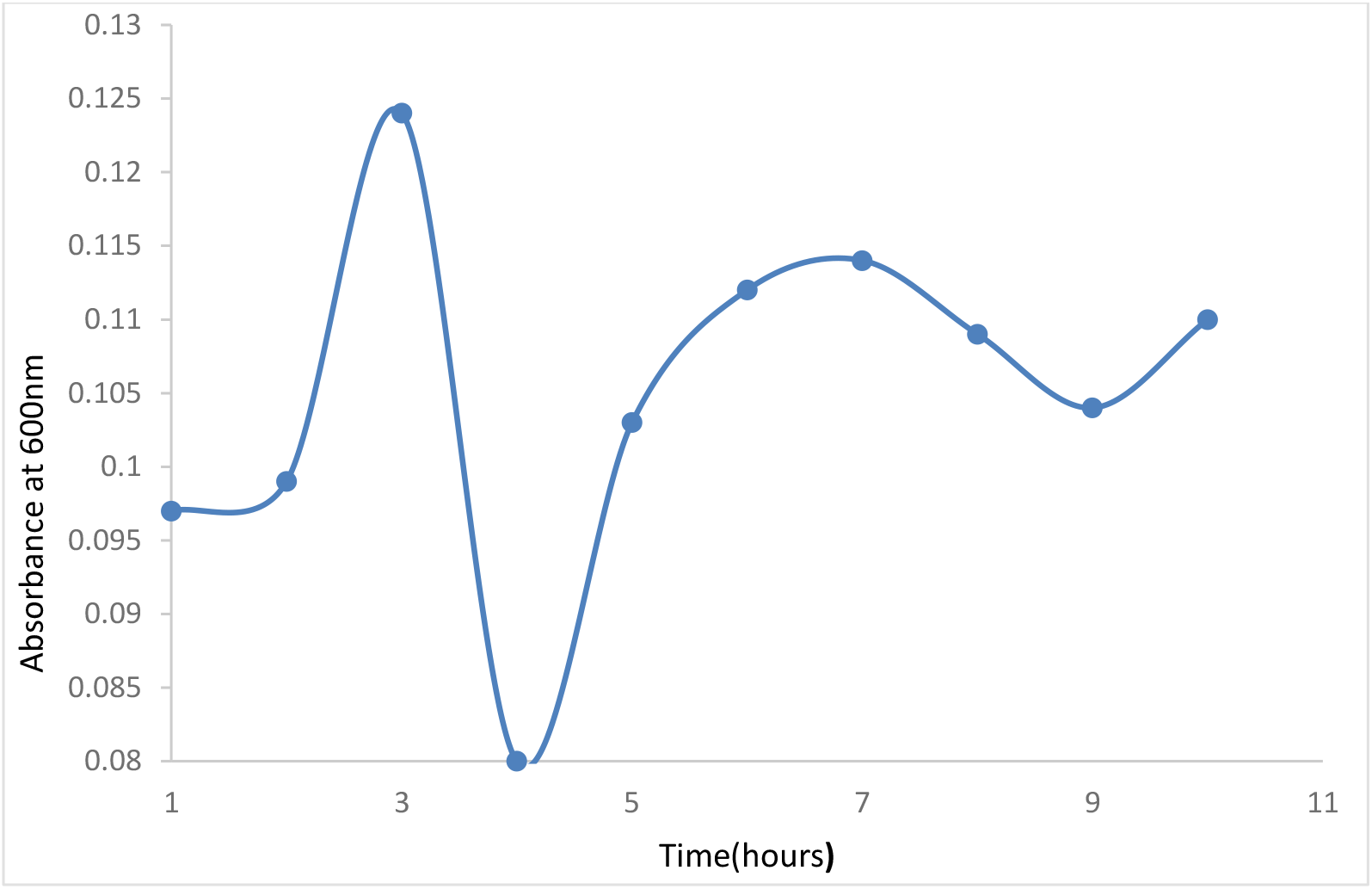
Growth Curve of *P. aeruginosa* for ten hours. The broth was kept at room temperature and absorbance read at an hourly interval also for ten hours at 600nm. The experiment was conducted in triplicate.

### Level of Biofilm Formed by *Pseudomonas aeruginosa* in the two different catheters

The level of biofilm formed by *P. aeruginosa* in the two catheters is shown in Figure 2. After the first day, the level of biofilm formed in the silicon catheter (0.064) was higher than that of the latex coated (0.053). But after the second day and third day the level of biofilm formed in the latex coated catheter (0.138 and 0.157 for day 2 and 3 respectively) were higher than that of the silicon (0.081 and 0.131 for day 2 and 3 respectively).

**Figure 2:**
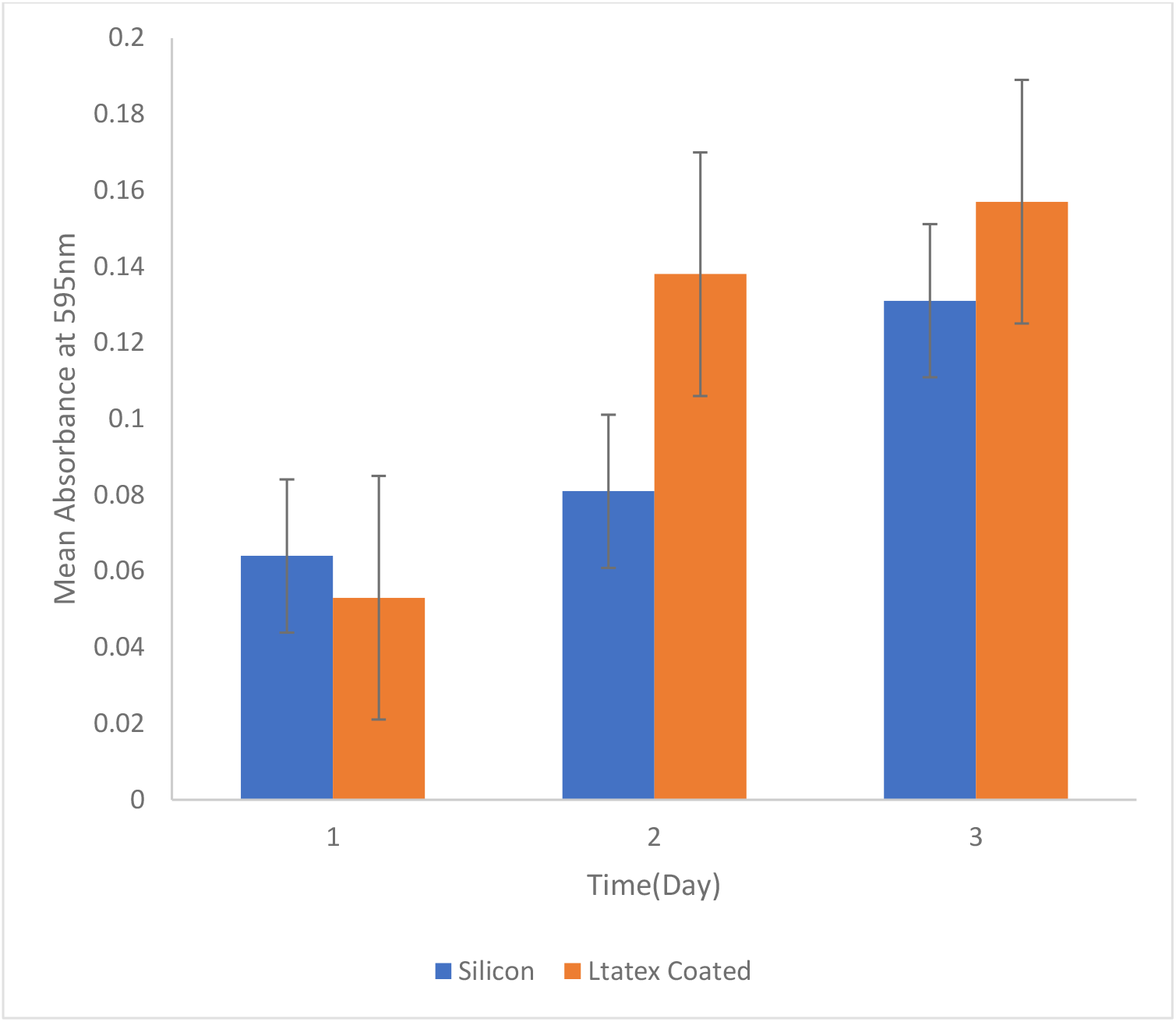
Level of biofilm formed by P *.aeruginosa* in two different catheters with respect to time

### Level of Biofilm Formed by *Pseudomonas aeruginosa* in Microplates

The level of biofilm formed of P. aeruginosa in microplates is shown in Figure 3. There were increases in level of biofilm as the days moved along (0.081, 0.085 and 0.103 for the first, second and third day respectively).

**Figure 3:**
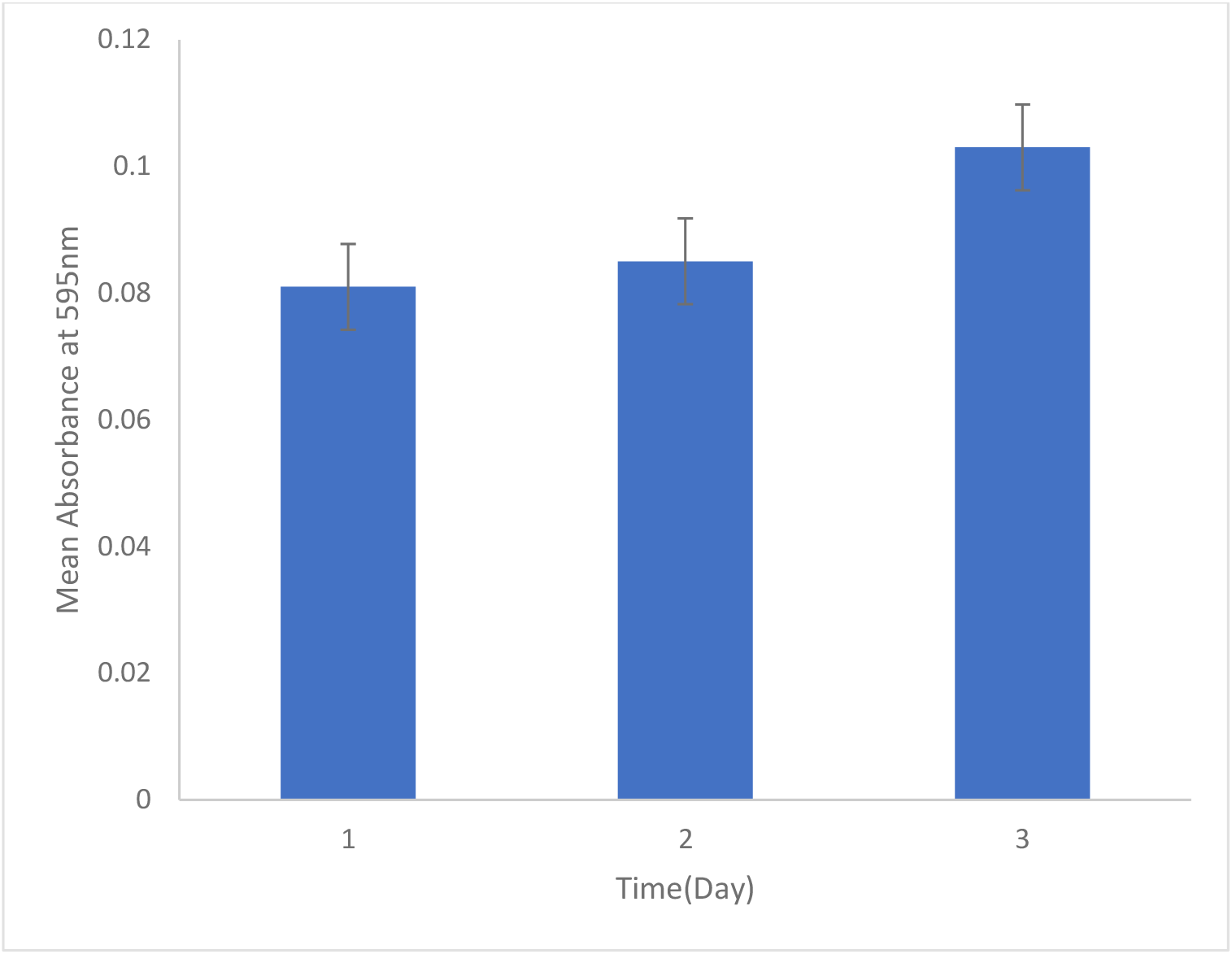
Level of biofilm formed by P.aeruginosa in microplates with respect to time

## DISCUSSION

Biofilm formation in urinary catheters is the major cause of urinary tract infections. There are different types catheters based on the material used in their manufacturing; the degree and rapidity of biofilm formation on a catheter is influenced by the material and chemicals used in manufacturing the catheter (Vaterrodt *et al.*, 2016). In contrast different types of catheters have different susceptibility levels for the formation of biofilm. The rate of biofilm formation in a catheter is also dependent of the type of bacteria involved (Dang and Lovell, 2016). It is therefore important to determine the growth curve of the bacterial which form the biofilm. As shown in figure 1, the growth curve of *P. aeruginosa* deviated from normal growth curve that was expected. This might be as a result of an error incurred during the experiment or might be a fault of the spectrophotometer used for measuring the absorbance.

### Level of Biofilm Formed in the Catheters

As shown in figure 2, the level of biofilm formed in the silicon catheter (0.064) was higher than that in the Latex coated catheter (0.053) after the first day. This might imply that, the rate of biofilm formation in silicon catheter is higher than that of Latex coated catheter. After the second day, the level of biofilm formed in the latex coated catheter (0.138) was far higher than that of the silicon catheter (0.081). And after the third day the level of biofilm formed in the latex coated catheter (0.157) was still higher than that of the silicon catheter (0.131). This suggests that the silicon catheter is less susceptible to *P. aeruginosa* for biofilm formation compared to the latex coated catheter therefore the need to include the assessment of biofilm in the quality evaluation of catheters. The observation after the first day might be as a result an experimental error or might suggest that materials that are less susceptible to bacteria for biofilm formation allow the formation of biofilm at a higher rate during the initial stages before showing a lower of growth. To justify this assumption, the formation of biofilm by the bacteria on another material was required.

Comparing the level of biofilm formed by the same bacteria *(P. aeruginos*a) in microplates to the level in the catheters, it was observed that the level of biofilm formed in the microplate for the first day (0.081) was higher than that of the catheters. The level of biofilm formed in the microplate for the second (0.085) and the third day (0.103) was lower compared to the catheters. This justified the earlier assumption that materials with lower susceptibility to *P. aeruginosa* for biofilm formation show a higher growth rate at the initial stage before showing a lower rate of biofilm formation (Hahnel *et al.*, 2015) and that the observation in the catheters after first day was not as a result of an experimental error. This might be for only *P. aeruginosa* and might not be seen when a different bacterium is used. This is because biofilm formation is dependent on the type of bacteria and the surface involved. It could also be realised that the level of biofilm formed in the microplate was lower than that of the urinary catheters which are more exposed to microbes when inserted into the bladder.The difference in the level of biofilm formed in the two catheters suggest that, there is the need to include the assessment of biofilm formation in the quality evaluation of catheters to help minimise the rate of urinary tract infections that are associated with the use of catheters. The formation of biofilms is regulated by mechanisms such as quorum sensing that help to strengthen and protect the cells. (Clayton *et al.*, 2017). Hence chemical modification that can alter these mechanisms can be considered during the manufacture and evaluation of urinary catheters.

In conclusion, latex coated catheter is more susceptible to *P. aeruginosa* for the formation of biofilm than silicone catheters. It is recommended that in the quality evaluation of catheters, the assessment of biofilm formation should be included. Also further studies should be done to investigate how different catheter material support biofilm formation and the mechanisms involved.

## ACKNOWLEDGEMENTS

This work was supported by the Department of Biochemistry, University of Cape Coast. The technical support from Eric Ofori (Analytical Laboratory) is acknowledged.

